# IRE1α regulates macrophage polarization, PD-L1 expression and tumor survival

**DOI:** 10.1101/2020.02.17.952457

**Authors:** Alyssa Batista, Jeffrey J. Rodvold, Su Xian, Stephen Searles, Alyssa Lew, Takao Iwawaki, Gonzalo Almanza, T. Cameron Waller, Jonathan Lin, Kristen Jepsen, Hannah Carter, Maurizio Zanetti

## Abstract

In the tumor microenvironment local immune dysregulation is driven in part by macrophages and dendritic cells that are polarized to a mixed proinflammatory/immune suppressive phenotype. The unfolded protein response (UPR) is emerging as the possible origin of these events. Here we report that the inositol-requiring enzyme 1 (IRE1α) branch of the UPR is directly involved in the polarization of macrophages *in vitro* and *in vivo,* including the upregulation of IL-6, IL-23, Arginase1, as well as surface expression of CD86 and PD-L1. Macrophages in which the IRE1α/Xbp1 axis is blocked pharmacologically or deleted genetically have significantly reduced polarization, and CD86 and PD-L1 expression, which was induced independent of IFNγ signaling suggesting a novel mechanism in PD-L1 regulation in macrophages. Mice with IRE1α- but not Xbp1-deficient macrophages showed greater survival than controls when implanted with B16.F10 melanoma cells. Remarkably, we found a significant association between the IRE1α gene signature and *CD274* gene expression in tumor-infiltrating macrophages in humans. RNASeq analysis showed that bone marrow derived macrophages with IRE1α deletion lose the integrity of the gene connectivity characteristic of regulated IRE1α-dependent decay (RIDD) and the ability to activate *CD274* gene expression. Thus, the IRE1α/Xbp1 axis drives the polarization of macrophages in the tumor microenvironment initiating a complex immune dysregulation leading to failure of local immune surveillance.

## INTRODUCTION

Myeloid cells in the tumor microenvironment (TME) are of central relevance to understand the dynamics of tumor progression [1]. They infiltrate tumors in varying numbers depending on tumor types and display phenotypic and functional diversity [2, 3]. Among them macrophages and dendritic cells-cells privileged with antigen presentation/T cell activation functions-often acquire a mixed pro-<u>i</u>nflammatory/<u>i</u>mmune <u>s</u>uppressive (IIS) phenotype, both in the mouse [4, 5] and in humans [6, 7]. Because this phenomenon is considered at the root of the dysregulation of local adaptive T cell immunity [8, 9], much emphasis has been placed on identifying common mechanisms driving the acquisition of tumor-promoting properties by macrophages and dendritic cells in the TME [5, 10–14].

The TME is home to environmental *noxae* such as hypoxia and nutrient deprivation [15]. In addition, about 20% of tumors have a viral origin [16] and most (90%) solid tumors carry chromosomal abnormalities [17]. These events, independently or collectively, can lead to a dysregulation of protein synthesis, folding, and secretion [18, 19], and the accumulation of misfolded proteins within the endoplasmic reticulum (ER), triggering a stress response termed the unfolded protein response (UPR) [20]. The UPR, an evolutionarily-conserved adaptive mechanism [21], is mediated by three initiator/sensor ER transmembrane molecules: inositol-requiring enzyme 1 (IRE1α), PKR-like ER kinase (PERK), and activating transcription factor 6 (ATF6). In the unstressed state these three sensors are maintained inactive through association with the 78-kDa glucose-regulated protein (GRP78) [22]. During ER stress, GRP78 disassociates from each of the three sensors to preferentially bind un/misfolded proteins, activating each sensor and their downstream signaling cascades, which aim to normalize protein folding and secretion. PERK, a kinase, phosphorylates the translation initiation factor 2 (eIF2α) that effectively inhibits translation of most mRNAs, ultimately reducing ER client proteins. IRE1α, also a kinase, auto-phosphorylates and activates its RNase domain, resulting in the cleavage of the X-box binding protein 1 (XBP1) mRNA, yielding the production of the potent spliced XBP1 transcription factor isoform (XBP-1s), which drives the production of various ER chaperones to restore ER homeostasis. XBP-1s also binds to the promoter of several pro-inflammatory cytokine genes [23]. In addition, under ER stress or enforced autophosphorylation, IRE1α RNase domain can initiate an endonucleolytic decay of many ER-localized mRNAs, a phenomenon termed regulated IRE1α-dependent decay (RIDD) [24]. ATF6, a transcription factor, translocates to the Golgi where it is cleaved into its functional form, and acts in parallel with XBP-1s to restore ER homeostasis [25]. If ER stress persists despite these compensatory mechanisms, the transcription factor 4 (ATF4) downstream of eIF2α activates the transcription factor CCAAT-enhancer-binding protein homologous protein (CHOP) to initiate apoptosis [20].

Although the UPR serves essentially as a cell-autonomous process to restore proteostasis, it can also act in a cell-nonautonomous way through the release of soluble molecules, a phenomenon likely to occur when cancer cells undergo an acute or unresolvable UPR [26, 27]. Stress signals emanating from the ER of ER stressed cancer cells may thus result in the induction of ER stress in neighboring cells, including macrophages and dendritic cells [26, 28]. This sets in motion a broad range of adaptive responses creating a functional cooperation or community effect among cells in the TME [29–31]. Under controlled experimental conditions bone marrow-derived macrophages and dendritic cells (BMDM and BMDC) cultured in conditioned media of ER stressed cancer cells develop a de novo UPR and acquire a mixed IIS phenotype [26, 28] characterized by the transcriptional upregulation of the tumorigenic pro-inflammatory cytokines IL-6, TNFα, and IL-23 [32–34], and contextually of the immune-suppressive enzyme Arginase 1 (Arg1) [35]. Under these conditions, cross-priming of naïve CD8^+^ T cells by BMDC is greatly compromised [28]. In line with this observation, Cubillos-Ruiz reported that the incubation of BMDC in ovarian cancer conditioned media results in *Xbp1* splicing, and that the conditional knock-out of *Xbp1* in dendritic cells improves antigen presentation and significantly reduces tumor growth *in vivo* [36]. In line with these observation is a report showing that GRP78 in cancer cells regulates macrophage recruitment to mammary tumors through metabolites secreted from cancer epithelial cells [37]. Thus, UPR-driven cell-nonautonomous mechanisms play a hitherto unappreciated role in orchestrating immune cells in the TME and driving their dysregulation, so as setting the stage for failure of local immune surveillance.

We therefore decided to elucidate the mechanism(s) through which the UPR may ultimately affect immune cells and perturb the TME to promote tumor growth. We focused on macrophages as these cells represent the major population infiltrating most solid tumors in humans, conspicuously more abundant than dendritic cells and other cells of myeloid origin [38]. Relative to dendritic cells or myeloid derived suppressor cells (MDSCs) [39, 40] little is known about how the UPR affects macrophages during cancer development. Based on our earlier report that BMDM can be polarized to a mixed IIS phenotype via a UPR-mediated cell-nonautonomous mechanism [26] our initial goal was to verify whether this phenomenon could be recapitulated in tumor-infiltrating macrophages *in vivo* in immunocompetent mice, and what UPR pathway might contribute to their dysregulation. To this day, these questions have remained largely unanswered. Here we show that the UPR and the IRE1α/XBP1 axis are activated in macrophages during tumor growth, that the conditional knock-out of IRE1α in macrophages regulates the acquisition of a mixed IIS phenotype and is also sufficient to restrain tumor development *in vivo*. Importantly, we discovered that IRE1α signaling regulates PD-L1 expression in murine and in tumor-infiltrating macrophages in humans.

## RESULTS

### Tumor infiltrating CD11b^+^ myeloid cells display the UPR/IIS signature *in vivo*

Previous *in vitro* studies indicated that BMDC and BMDM respond to a cell-nonautonomous UPR developing a complex phenotype characterized by a UPR activation and a mixed pro-inflammatory/immune suppressive (IIS) phenotype [26, 28]. Here as an initial step we interrogated tumor-infiltrating myeloid cells (CD11b^+^) to document these characteristics during tumor growth *in vivo*. To this end, we implanted B16.F10 murine melanoma cells into C57BL/6 mice that carry the *Xbp1-Venus* fusion transgene under the control of the CMV-β actin promoter, known as the ER stress-activated indicator (ERAI) [41], which reports IRE1α mediated *XBP1* splicing through the expression of the fluorescent Venus protein. First, we interrogated the relative abundance of CD11b^+^ cell infiltrate into tumors three weeks after implantation of B16.F10 tumor cells and found that 2-5 % of the bulk tumor consisted of CD11b^+^ myeloid cells (Fig. S1). Of these ∼50% expressed the F4/80 surface marker specific of macrophages. We then compared the expression of the *Venus* protein in tumor-infiltrating CD11b^+^ cells to those in the spleen and bone marrow, both from tumor-distal and tumor-proximal femurs (Fig 1A). The *Venus* protein signal was significantly higher in tumor-infiltrating CD11b^+^ cells relative to those in control tissues, suggesting a concurrent UPR signaling with *XBP1* splicing in the TME only.

**Figure 1.**
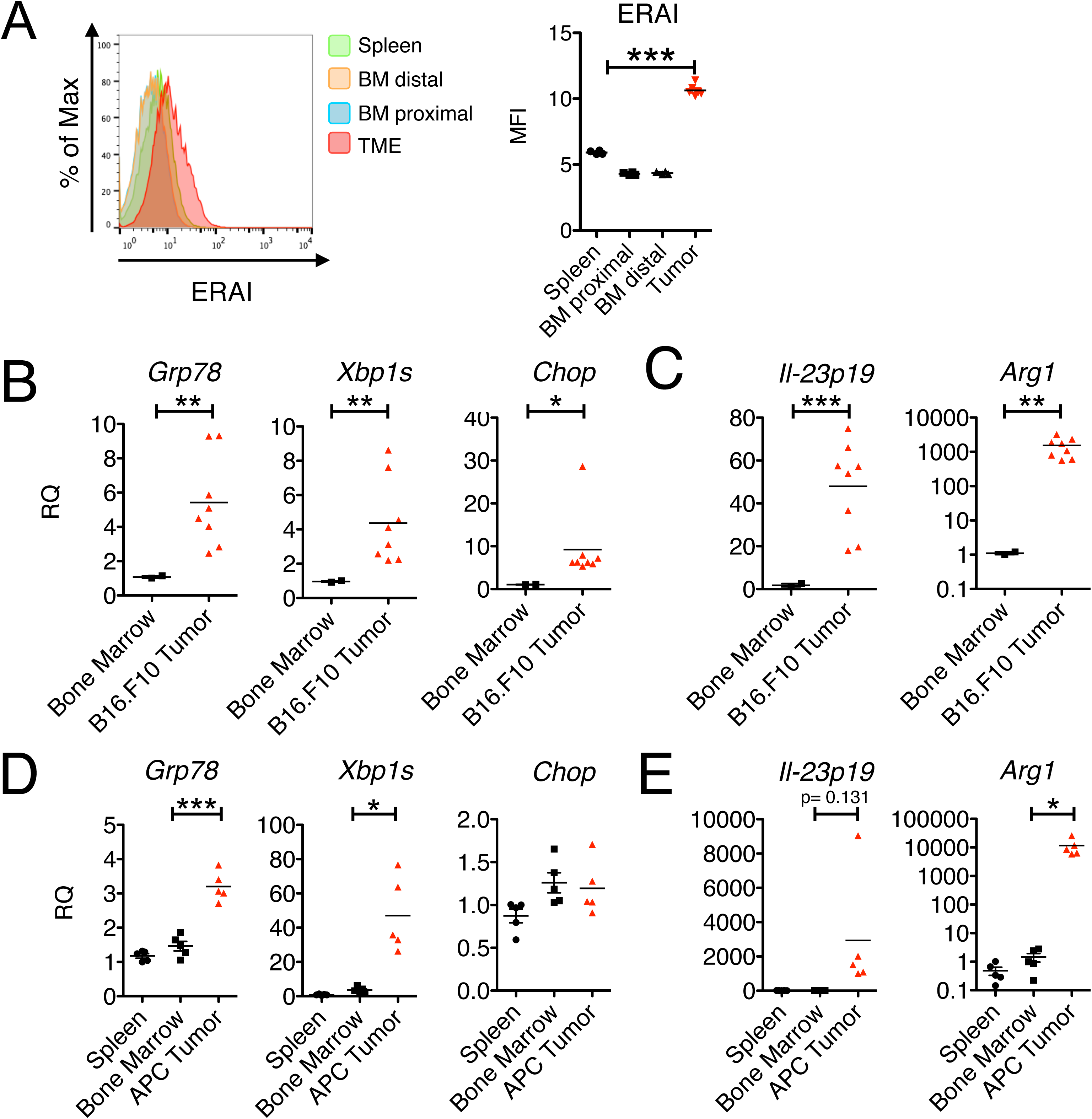
Activation of the UPR and acquisition of the IIS phenotype by tumor-infiltrating CD11b^+^ cells *in vivo*. (A) Flow cytometry histogram and comparative mean fluorescent intensity (MFI) values (n=4) of ERAI expression in CD11b^+^ cells resident in specified tissue. (B,C) Gene expression in CD11b^+^ cells isolated from B16.F10 tumors and respective bone marrow (n≥2 /group). Gene expression was arbitrarily normalized to one bone marrow sample and values represent relative quantification (RQ) fold transcription expression. (D,E) Gene expression in CD11b^+^ cells isolated from APC adenomas, and respective bone marrow and spleen (n≥2 / group). RNA extracted from these cells was analyzed by RT-qPCR using specific primers.

Having established that *XBP1* splicing occurs in tumor-infiltrating CD11b^+^ cells, we sought to detect other features of the IIS phenotype. To this end, we implanted B16.F10 cells in wild-type C57BL/6 mice and isolated by positive selection CD11b^+^ cells from tumor, spleen and bone marrow 22 days post-implantation. Phenotypically, the isolated cells were CD11b^+^ and Gr1^-^ and showed the transcriptional upregulation of three key UPR genes: *Grp78*, a downstream target of the ATF6 pathway, spliced *Xbp1* (*Xbp1-s*) a downstream product of the IRE1α pathway, and *Chop*, a downstream product of the PERK pathway (Fig. 1B). A transcriptional upregulation of all three genes suggested the activation of a classical UPR. Contextually, CD11b^+^ cells also showed the transcriptional upregulation of *Il23p19,* a key pro-inflammatory cytokine gene, and Arginase-1 (*Arg1*), an immune suppressive enzyme (Fig. 1C).

To see if the UPR/IIS signature also hallmarks CD11b^+^ cells during spontaneous tumor growth, we interrogated mice with mutations in the adenomatous polyposis coli (*Apc*) gene (“Apc mice”), which develop small intestinal adenomas by 30 days of age [42]. We pooled CD11b^+^ cell infiltrates from adenomas from multiple Apc mice and probed the expression of UPR genes, *Il-23p19* and *Arg1* relative to CD11b^+^ cells isolated from either the bone marrow or the spleen as controls. CD11b^+^ cells from APC adenomas had increased expression of UPR genes, *Il-23p19* and *Arg1* (Fig. 1D,E). Collectively, these data suggest that CD11b^+^ cells infiltrating the TME undergo ER stress and are polarized to the IIS phenotype.

### IRE1α dependent cell-nonautonomous polarization of macrophages

Environmental conditions shown to have tumor promoting effects have been linked to both IRE1α and PERK, making it necessary to determine which of the two was responsible for the acquisition of the IIS phenotype in our model system. To probe the role of IRE1α, we used the small molecule 4μ8C, an inhibitor specific for the RNAse domain. This small molecule forms an unusually unstable Schiff base at lysine 907 (K907) and inhibits both XBP1 splicing and regulated IRE1α-<u>d</u>ependent <u>d</u>ecay (RIDD), but not IRE1α kinase activity. To confirm that 4µ8c (30 µM) was effective we measured *Xbp-1* splicing in C57Bl/6 BMDM and B16.F10 cells treated with the conditioned medium (CM) of ER stressed cancer cells (transmissible ER stress conditioned medium or “TERS CM”) (Fig. S2). Compared to uninhibited conditions, 4μ8C did not significantly affect the transcriptional of UPR genes (*Grp78* and *Chop*, Fig. 2A). However, it significantly inhibited the transcriptional activation of *Il-6* and *Il-23p19* (Fig. 2B) and trended towards inhibiting *Arg1* (p=0.127) (Fig. 2C). Previously, we showed that TERS CM promotes the expression of CD86 and PD-L1 in BMDC [28]. Herein, we determined that ERAI BMDM treated with TERS CM also upregulate CD86 and PD-L, and that such an upregulation that is markedly inhibited by 4µ8C (Fig. 2D).

**Figure 2.**
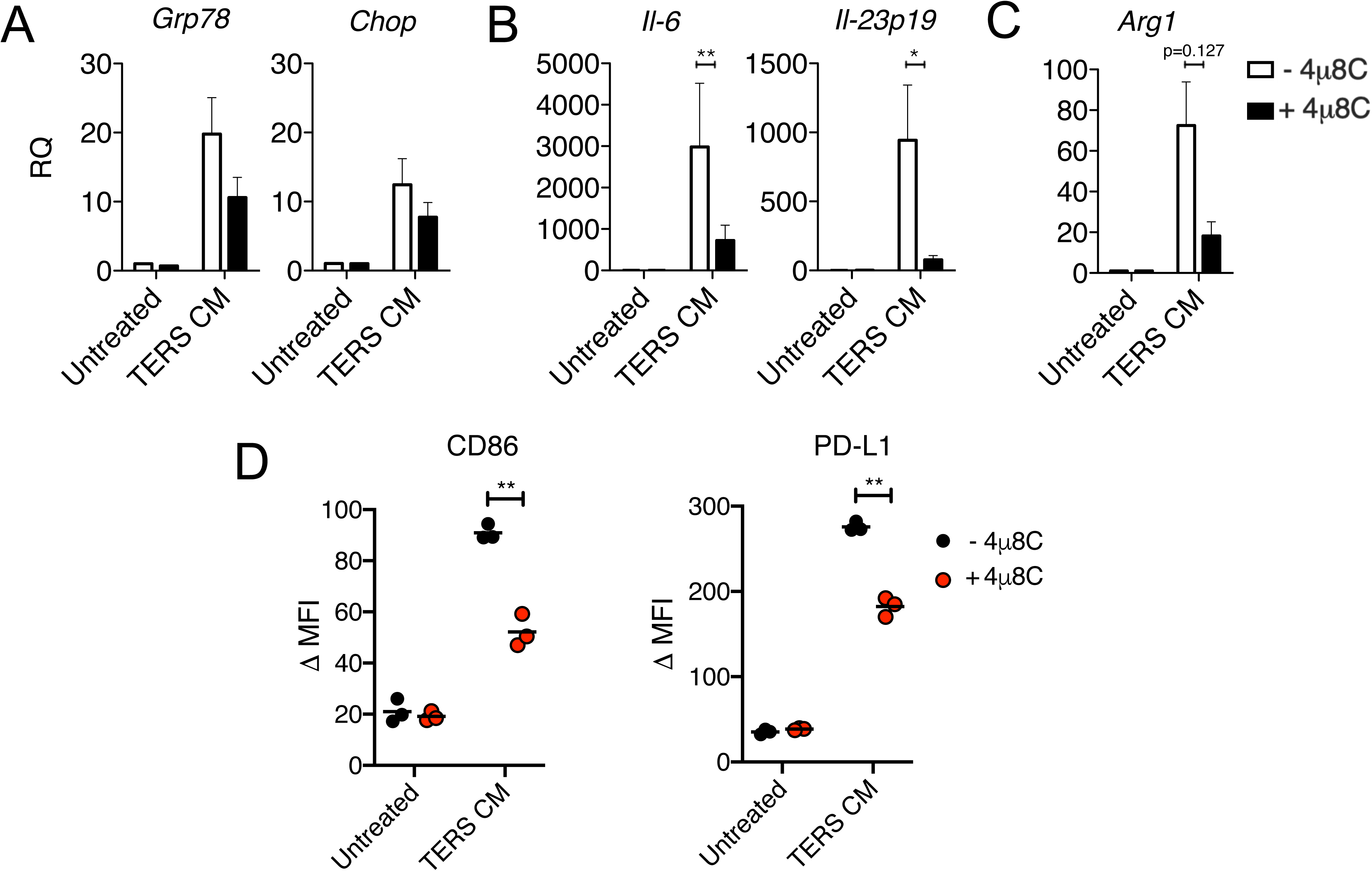
Chemical IRE1α inhibition prevents IIS polarization of BMDM *in vitro*. BMDM were culture *in vitro* in conditioned medium of ER stressed cancer cells (TERS CM) for 18 hours with or without 4u8C (30 μΜ) and their mRNA subsequently tested by RT-qPCR to detect the expression of (A) UPR genes (Grp78 and Chop) (B) pro-inflammatory cytokines (*Il6* and *Il23p19*), and (C) immune suppression genes (*Arg1*) (n=3-5/group). Relative quantification (RQ) was determined by arbitrarily normalizing gene expression to a Vehicle CM condition. Data points are expressed as means ±SEM. (D) Flow cytometry analysis of the intracellular expression of Venus protein (ERAI), and CD86 and PD-L1 surface expression in BMDM treated with conditioned medium of ER stressed tumor cells (TERS CM) with or without 4u8C (30 μΜ).

The involvement of the PERK pathway on the acquisition of the IIS phenotype by BMDM was assessed using the small molecule GSK2656157, a preferential PERK inhibitor [43]. GSK2656157 efficiently inhibited PERK phosphorylation (Fig S3A) but had no effect on the upregulation of *Grp78, Il-6* and *Arg1* induced in BMDM cultures by TERS CM (Fig. S3B). Congruently, PERK inhibition had little to no effect on the surface expression of CD86 and PD-L1 (Fig S3C). Collectively, these results suggest that BMDM polarization to the IIS phenotype is IRE1α dependent.

The role of IRE1α during macrophage activation by stimuli not obviously related to the UPR was tested in experiments in which BMDM were activated by LPS, a canonical activator of macrophages, or two metabolites shown to be relevant to the function of myeloid cells in the tumor microenvironment: lactic acid [10] and 4-hydroxynonenal (4-HNE), a products of lipid peroxidation [36]. While none of these molecules induced the transcriptional activation of *Gr78,* LPS consistently and readily induced *Il23p19* and *Il6* independent of IRE1α. Lactic acid induced *Arg1* only, and 4HNE had no effect on any of the target genes studied. Interestingly, 4µ8C reduced the induction of *Arg1* by both LPS and lactic acid, suggesting that the IRE1α may regulate the expression of this immune suppressive molecule outside the context of the UPR (Fig. S4).

### Loss of IRE1α−Xbp1 in macrophages attenuates the IIS phenotype, PD-L1 expression and tumor growth *in vivo*

Earlier reports showed that XBP1 is required for the development and survival of bone marrow derived DC [44], and that the deletion of XBP1 in lymphoid DC [40, 45] or in tumor-associated DC [36] improves antigen cross-priming and reduces tumor (ovarian) growth in the mouse. The role of the IRE1α/XBP1 axis in macrophage activation in the context of tumorigenesis has not been previously explored. Chemical inhibition of IRE1α endonuclease activity clearly implicated the IRE1α pathway in macrophage polarization to the IIS phenotype. However, since 4µ8C inhibits both *Xbp1* splicing and RIDD activity [46], we used a genetic approach to distinguish mechanistically among the two IRE1α functions in the acquisition of the IIS phenotype. To this end, we developed mice with *Ern1* (the gene coding for IRE1α) or *Xbp1* conditional knockout (CKO) in macrophages by breeding mice floxed (*fl/fl)* for *Ern1* [41] or *Xbp1* [47] with LysM-Cre mice (B6.129P2-Lys2tm1(cre)Ifo/J [48]. The genotype of CKO mice is shown in Fig. S5. Western blot analysis of *Ern1* CKO BMDM confirmed the absence of IRE1α (Fig. 3A) as well as the absence of the spliced form of *Xbp1* following treatment with the SERCA (sarco/endoplasmic reticulum Ca^2+^-ATPase) inhibitor thapsigargin (Fig. 3B). Under similar experimental conditions, *Xbp1* CKO BMDM showed an intact IRE1α expression under basal conditions (Fig. 3A) but the absence of the spliced form of *Xbp1* after thapsigargin treatment (Fig. 3C). Thus, the LysM-Cre CKO system was effective at specifically deleting IRE1α and Xbp1 in activated BMDM.

**Figure 3.**
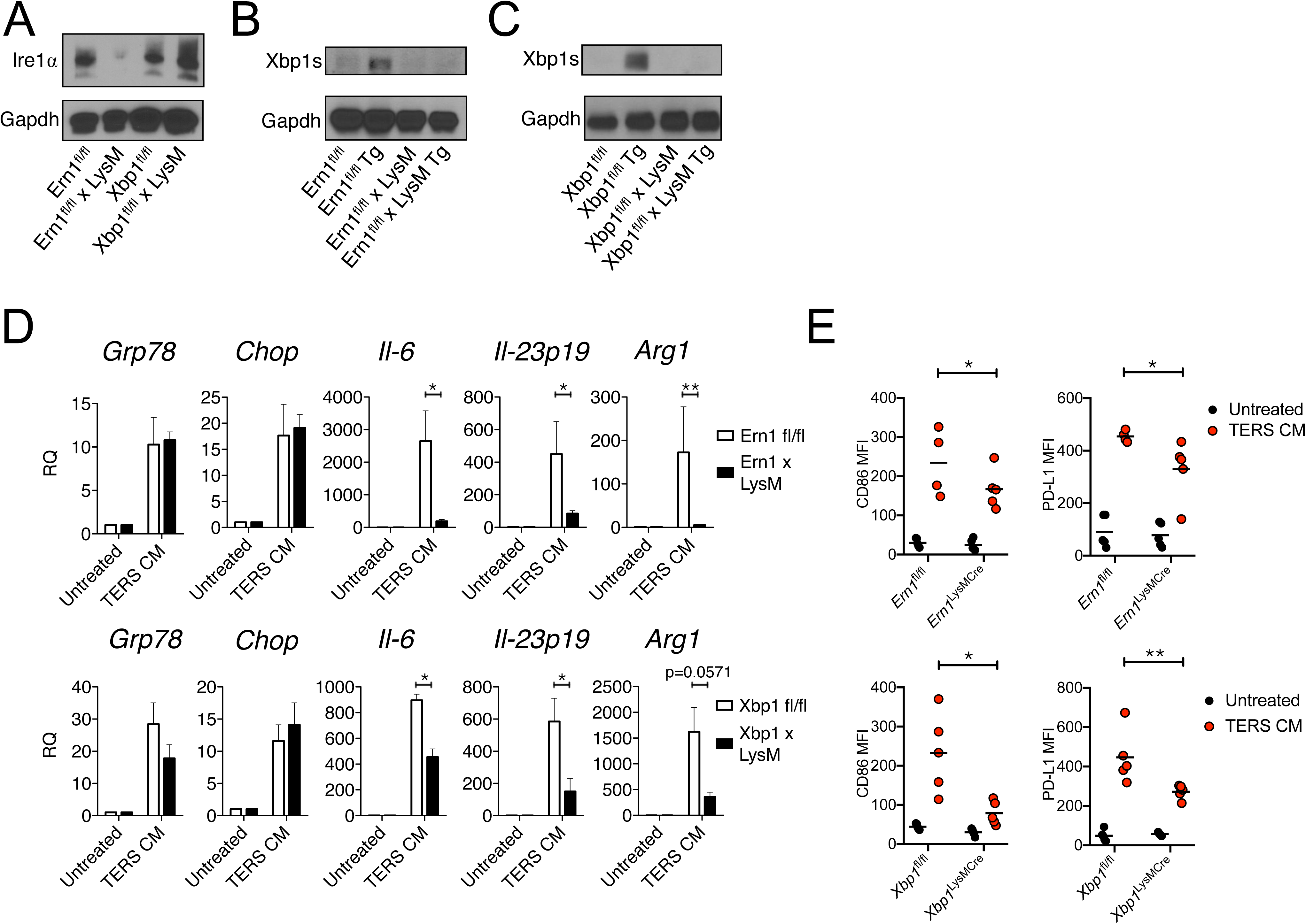
Deficiency in the IRE1α-XBP1 axis in macrophages attenuates the IIS phenotype, PD-L1 expression and tumor growth. (A) Western blot analysis of *Ern1* CKO BMDM showing lack Ire1upon activation (24 hrs) by thapsigargin (Tg) (300 nM). (B) Western blot analysis of *Ern1* CKO BMDM showing lack of spliced Xbp1 (Xbp1s) following activation (24 hrs) by by thapsigargin (Tg) (300 nM). (C) Western blot analysis of *Xbp1* CKO BMDM showing lack of spliced Xbp1 (Xbp1s) following activation (24 hrs) by thapsigargin (Tg) (300 nM). (D) RT-qPCR analysis of UPR and IIS genes in wild type or CKO BMDM untreated or treated with TERS CM. Values represent the mean ± SEM (n= 3-5/group). (E) IRE1-XBP1 deficiency reduces CD86 and PD-L1 expression in BMDM. *Ern1*fl/fl, *Xbp1*fl/fl, *Ern1* CKO and *Xbp1* CKO BMDM were treated (18 hrs) with TERS CM and subsequently stained with PE-conjugated antibodies to CD86 and CD274. The MFI for both surface proteins was quantified and plotted against the MFI of the corresponding unstimulated control. Statistical significance was determined using the Mann-Whitney *t* test. (n=4-5 mice/group).

First we compared the transcriptional response of *Ern1* and *Xbp1* CKO vs wild type BMDM when treated with TERS CM. We found that *Grp78* and *Chop* were unaffected in *Ern1* CKO BMDM, but *Il6*, *Il-23p19* and *Arg1* were markedly and significantly reduced in CKO relative to *fl/fl* control BMDM (Fig. 3D, upper panels). Likewise, in *Xbp1* CKO BMDM, the induction of *Grp78* and *Chop* was unaffected, but the activation of *Il6* and *Il23p19* was significantly reduced compared to *fl/fl* control BMDM. The activation of *Arg1* trended lower in *Xbp1* CKO compared to *fl/fl* control BMDM (p = 0.0571). (Fig. 3D, lower panels). These results confirm that the IRE1α-XBP1 axis mediates the IIS phenotype.

We then evaluated the effect of TERS CM on the expression of CD86 and PD-L1 in BMDM populations. *In vitro* treatment of *Ern1* or *Xbp1* CKO BMDM with TERS CM yielded a significant reduction of both surface proteins compared to wild type BMDM (Fig. 3E). Thus, the conditional deletion of the IRE1α/XBP1 axis in macrophages produced effects consistent with the pharmacological inhibition by 4µ8C. This suggests that the IRE1α-XBP1 axis is central to both macrophage activation (CD86 upregulation) and the acquisition of PD-L1, a marker of immune disfunction. We ruled out the possibility that PD-L1 expression was the result of canonical IFN-γ signaling since (a) we did not detect IFN-γ in TERS CM (Fig. S6A), (b) a blocking antibody to human IFN-γ had no effect on *Cd274* gene expression in BMDM treated with TERS CM (Fig. S6B), and (c) RNASeq data showed no induction of the *Ifng* gene in either *Ern1* CKO or *fl/fl* control BMDM treated with TERS CM (Fig. S6C).

To ascertain the physiological relevance of these findings, we next assessed the survival of *Ern1* and *Xbp1* CKO mice implanted with B16.F10 melanoma cells. We reasoned that survival would constitute an optimal initial read-out for the complex interactions between cancer cells and immune cells in the TME with focus on the IRE1α-XBP1 axis in myeloid cells. Survival in *Ern1* CKO mice was significantly greater (p=0.03) than in control *Ern1 fl/fl* mice (Fig. 4A). By contrast, *Xbp1* CKO mice survived longer than control *Xbp1 fl/fl* mice but the difference was non-significant (Fig. 4A). Based on survival data we isolated F4/80 tumor-infiltrating macrophages of tumor-bearing *Ern1* CKO mice to assess the UPR/IIS and *Cd274* gene expression status. *Xbp1s*, *Il-23p19*, *Arg1 and Cd274 genes* were all markedly reduced in *Ern1* CKO macrophages compared to their *Ern1 fl/fl* counterpart (Fig. 4B). Together, these results point to macrophage IRE1α as a key negative regulator of TME immunodynamics and tumor growth *in vivo*.

**Figure 4.**
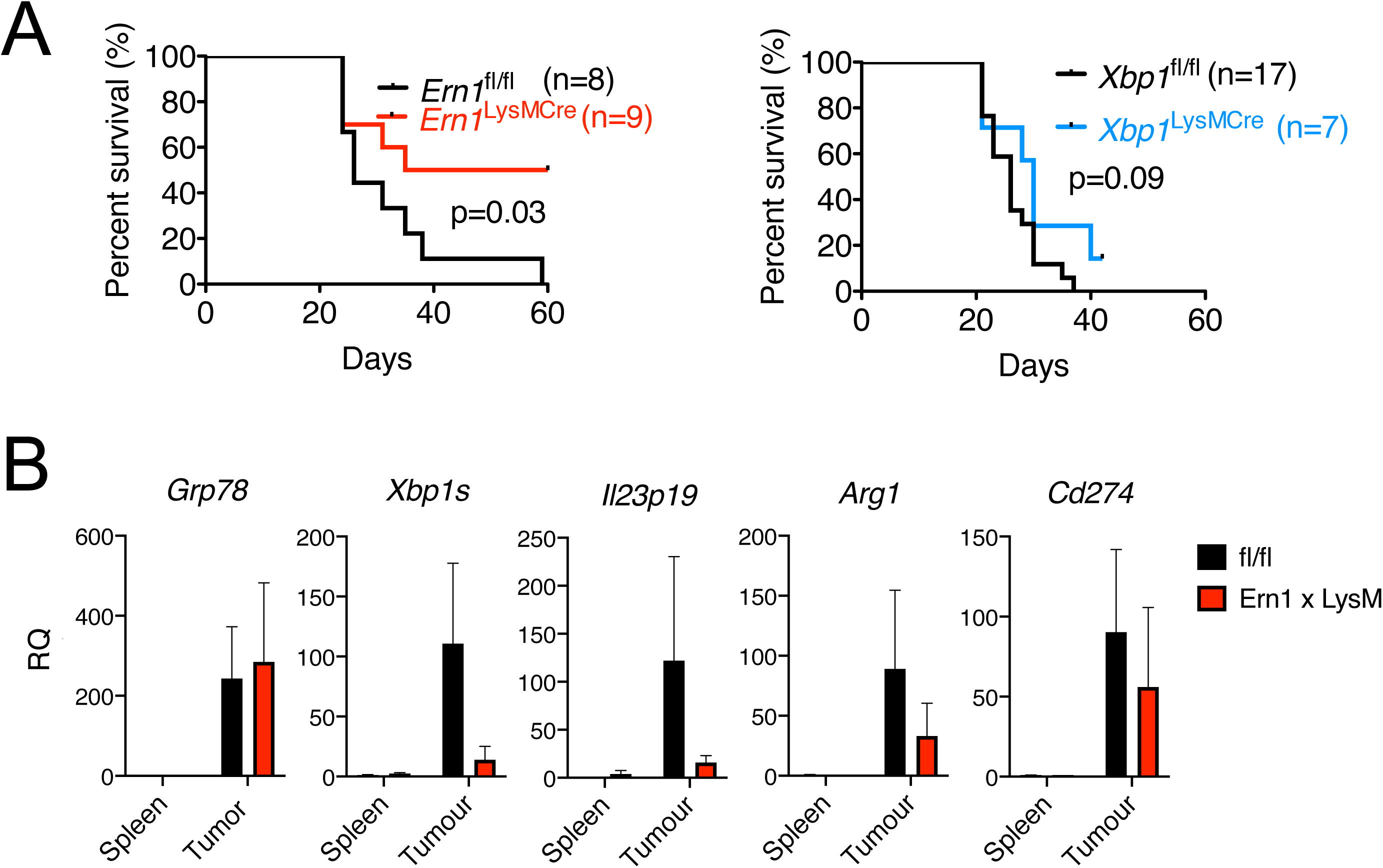
Tumor growth and tumor-infiltrating macrophage analysis in *Ern1/Xbp1* conditional knock out mice. (A) Kaplan-Meier survival curves of *Ern1fl/fl, Xbp1fl/fl, Ern1* CKO and *Xbp1*CKO mice injected in the right flank with 3×10e4 B16.ERAI cells/mouse. Tumor measurements were taken every two days in two dimensions. Mice were sacrificed once tumors reached 20 mm in either dimension. (B) Gene expression in F4/80^+^ macrophages isolated from B16.F10 tumors implanted in *Ern1* CKO or *fl/fl* mice, and respective spleen controls (n=2/group). mRNA was extracted enzymatically using the Zygem RNAgem Tissue PLUS kit. Gene expression was arbitrarily normalized to one spleen sample and values represent relative quantification fold transcript expression. Data points are expressed as means ±SEM.

### Loss of RIDD regulation in *Ern1* CKO macrophages

Because the IRE1α-XBP1 axis also regulates PD-L1 expression and both *Ern1* and *Xbp1* CKO BMDM showed significantly-reduced surface PD-L1 protein expression compared to *fl/fl* BMDM (Fig. 3E), we decided to distinguish the relative contribution of *Xbp1* splicing and RIDD to this phenomenon. To this end, we performed RT-qPCR on *Ern1-* and *Xbp1* CKO BMDM treated or not with TERS CM relative to *fl/fl* controls. We found that *Cd274* gene transcription was markedly and significantly lower in *Ern1* CKO BMDM relative to *fl/fl* controls (Fig. 5A). By contrast, *Xbp1* CKO BMDM and *fl/fl* BMDM had comparable *Cd274* gene transcription values (Fig. 5A). Based on this result and on PD-L1 surface expression (Fig. 3E), we tentatively conclude that XBP1-mediated regulation of PD-L1 occurs at the post-translation level, whereas IRE1α-mediated regulation is a transcriptional event. This conclusion favors the view that IRE1α-mediated PD-L1 regulation may occur via RIDD, justifying an in-depth analysis of RIDD activity in *Ern1* CKO BMDM.

**Figure 5.**
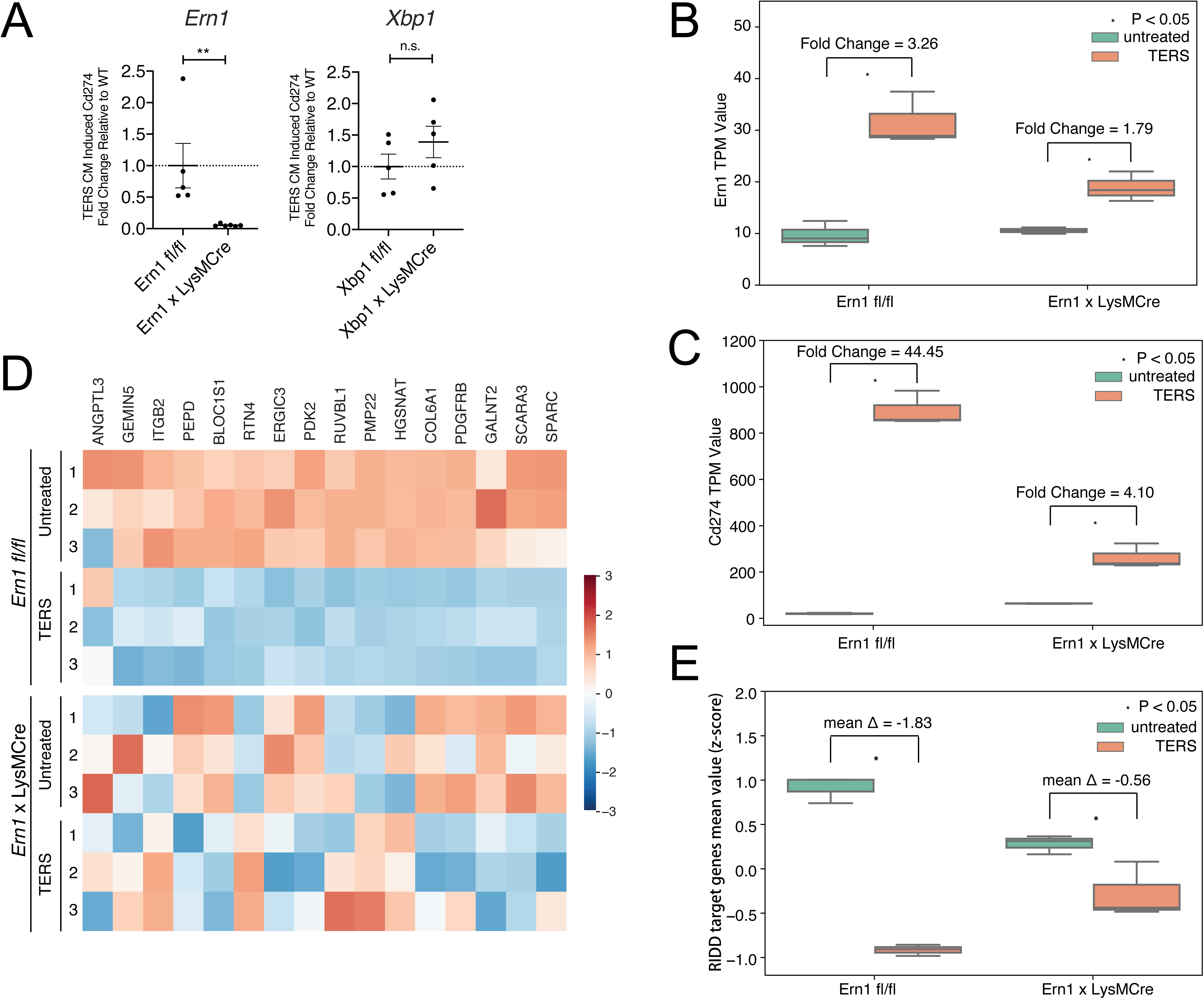
RIDD analysis of wild type and *Ern1* CKO BMDM treated with TERS CM. (A) Fold change in *Cd247* (PD-L1) transcription in *Ern1* deficient (left panel) and *XBP1* deficient (right panel) bone marrow-derived macrophages activated with TERS CM. (B) RNASeq analysis of *Ern1* expression in untreated or TERS CM treated wild type or *Ern1* CKO BMDM. TERS CM-induced fold changes are indicated in the graph. (C) Heatmap showing te relative expression of 16 RIDD target genes in untreated or TERS CM-treated wild type or *Ern1* CKO BMDM. (D) RNASeq analysis of *Cd274* expression in untreated or TERS CM treated wild type or *Ern1* CKO BMDM. TERS CM-induced fold changes are indicated in the graph. (E) Comparison of mean z-scores for the 16 RIDD target genes in untreated or TERS CM-treated wild type or *Ern1* CKO BMDM.

We performed RNASeq analysis of *fl/fl* and *Ern1* CKO BMDM untreated or treated with TERS CM. Three independently-derived BMDM populations per group were analyzed. The genotype of each mouse used in this experiment is shown in Fig. S5. Upon TERS CM treatment *Ern1* expression in *Ern1* CKO macrophages was 1.79-fold over that of untreated cells compared to 3.26 fold in *fl/fl* macrophages (Fig. 5B). We found that consistent with the flow cytometry data, *Cd274* (PD-L1) expression was markedly increased in macrophages (44.45-fold) but only moderately increased in *Ern1* CKO macrophages (4.11-fold, Fig 5C). Thus, both genetic and chemical inhibition of IRE1α signaling yielded concordant results.

Next, we performed a comprehensive analysis of RIDD activity using a set of 33 putative RIDD target genes previously defined [49]. We found that only half (sixteen) of these genes behaved as *bona fide* RIDD targets in TERS CM-treated BMDM (i.e., decreased expression after TERS CM treatment in *fl/fl* macrophages) (Fig. 5D, upper panel). We found that in *Ern1* CKO macrophages, there was a clear loss of a “RIDD signature” compared to *fl/fl* macrophages, both basally and after TERS CM treatment (Fig. 5D, lower panel). When considered together through an analysis of the mean z-score for the 16 genes, it became apparent that TERS CM induction of RIDD activity was much more effective in *fl/fl* than in *Ern1* CKO macrophages (Fig. 5E). Collectively, these results show that macrophages lacking *Ern1* lose RIDD regulation, suggesting that RIDD may be implicated in the regulation of PD-L1 expression.

In the same analysis we found that *Tabpb* (tapasin), a chaperone molecule involved in the stabilization of high affinity peptide/MHC-I complexes in the endoplasmic reticulum [50], did not behave as RIDD. In fact, *fl/fl* macrophages treated with TERS CM showed *increased* not diminished expression at variance with previous reports on lymphoid (CD8α^+^) dendritic cells [40, 45]. The expression of *Bloc1s1* (a canonical RIDD target) was reduced, confirming that TERS CM induces RIDD (Fig. S7A). RT-qPCR analysis of *Tapbp* in *Xbp1 fl/fl* macrophages showed similar results (Fig. S7B). Perhaps, *Tapbp* is regulated by RIDD differently in CD8α^+^ dendritic cells and in BMDM.

### A link between IRE1α and PD-L1 expression in human tumor-infiltrating macrophages

The data reported herein suggest that *Cd274* gene expression in murine macrophages is positively regulated by IRE1α. Recently, Xu et al. [51] reported that PD-L1 protein expression in murine MYC^tg^:KRAS^G12D^ tumor cells is decreased by a small molecule that enables the cell to resume translation while the eIF2α downstream from PERK remains phosphorylated. Therefore, we decided to study the relationship between *CD274* (PD-L1) gene expression and the two major UPR pathways, IRE1α and PERK, across multiple human cancers. We began by interrogating the relative contribution of *ERN1* (IRE1α) and *EIF2AK3* (PERK) to *CD274* gene expression. In this analysis, we queried The Cancer Genome Atlas (TCGA) collection of RNA-sequencing expression data for bulk samples from thirty-one tumor types. Across these data, we observed that *ERN1* correlates strongly with *EIF2AK3* (Pearson correlation coefficient = 0.55; p < 1e-200) (Fig. S8A), and that both *ERN1* (p ≤ 1.46e-51) and *EIF2AK3* (p ≤ 1.62e-44) correlate positively with *CD274*, suggesting that the UPR plays a role in *CD274* gene expression. These correlations prompted us to further interrogate the relationship between *CD274*, *ERN1* and *EIF2AK3,* with respect to levels of infiltrating macrophages in bulk tumor samples approximated by a macrophage score derived from the geometric mean of three genes expressed by macrophages (*CD11b*, *CD68*, and *CD163)*. We found a positive correlation between *ERN1* and *CD274* within the high macrophage infiltration group (> 70th percentile) (Spearman correlation coefficient 0.18; p < 1.3e-21) (Fig. S8B). By contrast, the low macrophage infiltration group (< 30^th^ percentile) had a much weaker correlation (Spearman correlation coefficient 0.06; p < 0.001) (Fig. S8B). On the other hand, *EIF2AK3* and *CD274* within the high macrophage infiltration group (> 70th percentile) had a lower correlation (Spearman correlation coefficient 0.09; p < 1.9e-7) than in the corresponding *Ern1* group (Fig. S8C). Finally, *EIF2AK3* and *CD274* within the low macrophage infiltration group (< 30th percentile) had a surprisingly higher correlation (Spearman correlation coefficient 0.15; p < 8.32-15) than in the respective high macrophage infiltration group (Fig. S8C). Collectively, this analysis suggests that when macrophage infiltration is high, *Ern1* is a better predictor of *CD274* gene expression than *EIF2AK3*.

We also integrated the macrophage score with *ERN1* and *EIF2AK3* to predict *CD274* expression in an ordinary least squares (OLS) linear regression model, including the tumor type as a covariate (Table 1). We found that this model assigns significant, positive coefficients for the interaction terms of macrophages with *ERN1* (*ERN1*\*Macrophages, beta coefficient = 0.0012, p < 0.023) but not *EIF2AK3* (*EIF2AK3*\*Macrophages, beta coefficient = 0.0007, p < 0.155), suggesting that *ERN1* but not *EIF2AK3* is predictive of *CD274* gene expression within tumor-infiltrating macrophages in individual tumor types (Table 1). To validate these results we analyzed RNA-Seq data generated from macrophages isolated from thirteen patients with either endometrial or breast cancer [52]. We found a strong Pearson correlation coefficient between *ERN1* and *EIF2AK3* in these data (correlation coefficient 0.738; p < 0.003), suggesting UPR activation. Since IRE1α activity is a multistep and complex process [53] and may not be completely captured by *ERN1* expression levels, we derived a systemic representation of pathway activity controlled by IRE1α and by comparison PERK. We collected sets of downstream genes in the IRE1α and PERK pathways [54], and derived aggregate scores for each pathway from the mean expression signal of all detectable genes after z-score transformation. Since the transformed pathway scores could potentially amplify noise from genes with low expression, we applied filters to include only genes in each pathway with levels beyond a specific threshold (Fig. S9). We varied this filter threshold from zero to one thousand raw counts and then included the pathway activity scores in multiple OLS linear models to predict *CD274* across tumor-infiltrating macrophage samples (Fig. 6A). We found that a filter threshold of 100 counts effectively reduced noise while preserving signal from 84% of detectable genes in both the IRE1α and PERK pathways. In this model, the IRE1α score predicted *CD274* expression with a significant positive beta coefficient (beta coefficient = 21.043, p-value = 0.040), while the PERK score was non-significant (beta coefficient = 36.842, p-value = 0.103). This pattern of significant IRE1α coefficient and nonsignificant PERK coefficient was consistent across all filter thresholds (Fig. 6B). Comparing models wherein *CD274* expression was explained by IRE1α activity alone or by both IRE1α and PERK activity using the Aikake information criterion analysis shows that a model containing both is as 0.54 times as probable as the IRE1α alone to minimize the information loss (ΔAIC = 1.23). Taken together, these analyses suggest that the activation of *CD274* gene expression in tumor-infiltrating macrophages depends primarily on the IRE1α pathway.

**Figure 6.**
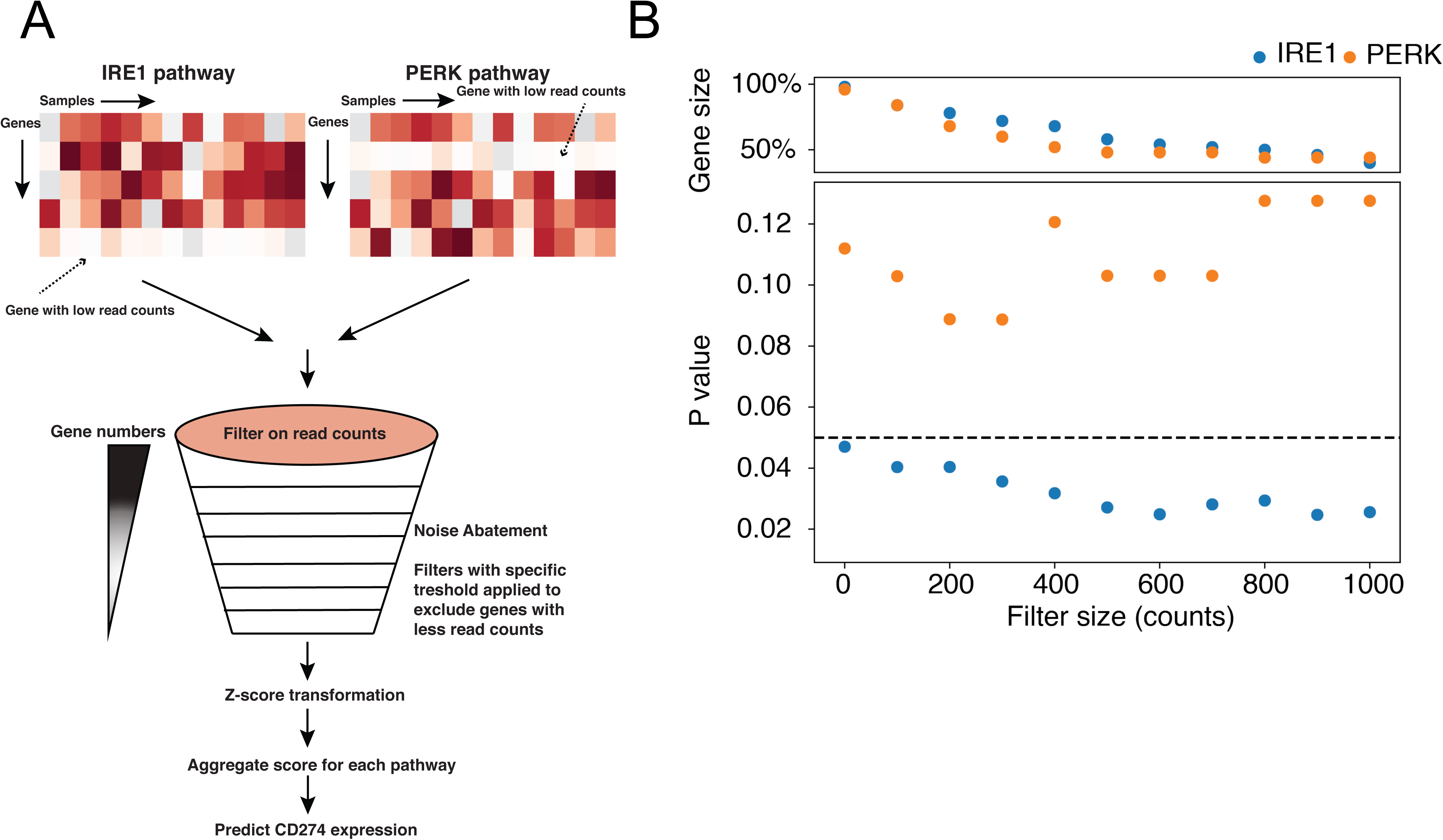
Ordinary least squares (OLS) linear model prediction of *CD274* gene expression in human tumor associated macrophages. (A) An illustration of the development of aggregated pathway scores. For both pathways we used filters with different thresholds to filter out genes with less read counts to account for baseline technical artifacts. Then we z-score transformed both the gene matrix for both pathways and aggregated these scores to predict *CD274* gene expression (B) RNAseq data from tumor associated macrophages isolated from 13 human endometrial or breast cancer samples were analyzed using 11 OLS linear models for each pathway (IRE1α or PERK). Each model was applied using different filters, each representing increasing read count thresholds. In the upper panel each dot represents the fraction of genes remaining in the model after a given filter was applied. In the lower panel the *p* value for each pathway predicting PD-L1 gene expression is indicated at each read count threshold.

Table 1.
|  | coef | pval |
| --- | --- | --- |
| Intercept | 0.5681224413 | 0.6307185602 |
| BLCA | 3.730781629 | 0.00291005966 |
| BRCA | -0.281147938 | 0.8147005789 |
| CESC | 6.893379114 | 8.34E-08 |
| CHOL | 0.4014684074 | 0.8413656853 |
| COAD | 0.5300701923 | 0.6672164335 |
| DLBC | 20.84235676 | 8.08E-29 |
| ESCA | 3.017860087 | 0.03778129109 |
| GBM | -0.5684440601 | 0.6883179244 |
| HNSC | 5.61635157 | 5.51E-06 |
| KICH | 5.178474502 | 0.0023769321 |
| KIRC | -0.3455225035 | 0.7796198465 |
| KIRP | 0.6659839637 | 0.6060961809 |
| LGG | -0.639698116 | 0.6071139362 |
| LIHC | -0.2908561171 | 0.8160364346 |
| LUAD | 3.69555473 | 0.00272696993 |
| LUSC | 6.865457981 | 3.49E-08 |
| MESO | 1.141219118 | 0.4692291251 |
| OV | -0.1591619966 | 0.9007001625 |
| PAAD | -0.7801816419 | 0.5683066888 |
| PCPG | 2.243029289 | 0.09976162006 |
| PRAD | -0.1264243114 | 0.9190474856 |
| READ | 0.3292890692 | 0.8124095707 |
| SARC | -1.090528135 | 0.4092896389 |
| SKCM | 0.8616842823 | 0.5776724443 |
| STAD | 4.070734989 | 0.001469900349 |
| TGCT | 0.872136964 | 0.5455941353 |
| THCA | 2.317361301 | 0.06056962238 |
| THYM | 13.04353041 | 4.96E-19 |
| UCEC | -0.04793209811 | 0.9687882727 |
| UCS | -0.5462166032 | 0.7546040883 |
| UVM | 0.1250809139 | 0.9379339045 |
| ERN1.00 | -0.02639975541 | 0.3447808132 |
| gmeanMacro | 0.04256029965 | 1.93E-07 |
| ERN1:gmeanMacro | 0.001211494277 | 0.02397204832 |
| EIF2AK3 | -0.002925408763 | 0.8832796928 |
| EIF2AK3:gmeanMacro | 0.0007820854511 | 0.1550669885 |

## DISCUSSION

Here we analyzed the effect of the UPR on gene expression regulation in macrophages as a potential mechanism driving immune dysregulation in the tumor microenvironment. Tumor-infiltrating CD11b^+^ myeloid cells in B16.F10 tumors and in spontaneously-arising colonic adenomas in Apc mice have an active UPR and display a mixed pro-inflammatory/immune suppressive phenotype. Using both a pharmacologic and genetic approach we show that the IRE1α/XBP1 axis plays a central role in macrophage activation and polarization to a mixed phenotype, including the upregulation of PD-L1. In agreement with the mouse data we found that in human tumor-infiltrating macrophages *CD274* (PD-L1) gene transcription correlates significantly with the IRE1α gene signature. B16.F10 tumor-bearing mice with conditional *Ern1*- but not *Xbp1* KO macrophages had significantly greater survival than their *fl/fl* controls. Collectively, these results show that IRE1α signaling drives macrophage dysregulation impacting negatively the immunobiology of the tumor microenvironment and ultimately the host’s ability to control tumor growth.

Virtually all adult solid tumors (carcinomas most notably) contain infiltrates of diverse leukocyte subsets, including macrophages, dendritic cells, and lymphocytes [2]. CIBERSORT and immunohistochemical tools have previously shown that macrophages represent the largest fraction among infiltrating leukocytes and their density correlates directly with poor survival [38, 55]. In the mouse, tumor-infiltrating CD11b^+^ myeloid cells produce pro-inflammatory/pro-tumorigenic cytokines (IL-6, IL-23, TNFα) [32–34], but oddly, also anti-inflammatory cytokines (IL-10, TGFβ) and molecules with immune suppressive function (Arginase1, perioxinitrite and indoleamine 2-3 dioxygenase) [8]. In humans, monocytes/macrophages with a “mixed” pro-inflammatory/suppressive phenotype have been reported in patients with renal cell carcinoma [6] and breast cancer [7]. Thus, a dysregulation-prone TME harbors CD11b^+^ myeloid cells with a split pro-inflammatory/immune suppressive phenotype that may be the result of hijacking by tumors for their own benefit [56]. Indeed, we previously proposed that tumor-derived UPR-driven factors determine the IIS phenotype in myeloid cells [57], contributing to progressive immune dysregulation and failure of immune surveillance.

Here, we analyzed two murine tumor models to demonstrate that tumor-infiltrating CD11b^+^ cells display features of UPR activation and a mixed IIS phenotype. The results clearly show that the UPR is associated with myeloid cell polarization *in vivo*, but do not allow a distinction between a cell-autonomous and a cell non-autonomous mechanism. However, since common triggers of inflammation such as LPS, or TME metabolites such as 4HNE and lactic acid [10, 36], did not induce a UPR/IIS phenotype, we favor the possibility that these changes in myeloid cells result from a cell non-autonomous mechanism of intercellular communication consistent with findings on BMDM and BMDC analyzed under controlled *in vitro* conditions [26, 28]. This appears to be a general mechanism since we recently showed that cell-nonautonomous intercellular communication among cancer cells induces an adaptive UPR imparting receiver cells with enhanced cellular fitness and resistance to various stressors [27].

A pharmacological approach using a small molecule (4µ8c) that inhibits IRE1α significantly reduced the transcription of *Il-6* and *Il-23p19* induced by TERS CM demonstrating a direct involvement of the IRE1α/XBP1 axis in driving pro-inflammation during an adaptive UPR. This is consistent with previous reports showing that XBP1 is recruited to the *Il6* and *Il23* promoters [23] and that *Il23* transcription is IRE1α-dependent [58]. Interestingly, 4µ8C did not reduce the transcription of these cytokines in the absence of a UPR, implying that IRE1α selectively regulates pro-inflammation within the boundaries of the UPR. Our findings on macrophage polarization via cell-nonautonomous means are consistent with reports showing that IRE1α drives M1 to M2 polarization of macrophages within white adipose tissue [59] and their inflammatory response to saturated fatty acids [60]. Importantly, 4µ8C also inhibited the TERS CM-induced upregulation of *Arg1,* and that of pro-angiogenic vascular endothelial growth factor (VEGF) (Fig. S10). Since IL-6 and IL-23 are known to bias T cell differentiation towards inflammatory (Th17) or regulatory T cells [61–65], and Arg1 potently suppresses the clonal expansion of T cells activated by antigen [28, 35], it follows that signaling through the IRE1α/XBP1 axis is of paramount importance to the economy of the TME and may be at the origin of a loss of local immune surveillance.

*Ern1* or *Xbp1* CKO macrophages enabled us to distinguish different roles within the IRE1α/XBP1 axis relative to immune dysregulation and tumor growth. *In vitro*, both *Ern1*- and *Xbp1*-CKO BMDM had decreased activation (CD86 and PD-L1 surface expression) and an attenuated IIS phenotype compared to control *fl/fl* macrophages when cultured in TERS CM, consistent with the effects of 4µ8C. However, only IRE1α deficiency significantly increased survival of mice implanted with B16.F10 melanoma cells, a result possibly reflected by an attenuation of the UPR/IIS signature and PD-L1 in tumor-infiltrating macrophages. Cubillos-Ruiz [36] also observed that IRE1α deficiency in DCs yielded greater survival than XBP1 deficiency in a model of ovarian cancer. By inference, we showed that B16.F10 tumor cells admixed with bone marrow-derived DCs with a UPR/IIS phenotype form faster-growing and larger tumors that had a marked reduction in tumor-infiltrating CD8^+^ T cells [28].

Chemical and genetic inhibition both showed that IRE1α regulates the surface expression of PD-L1 triggered in an IFNγ-independent manner through an adaptive UPR. PD-L1 activation is considered to occur mainly in response to IFNγ, albeit other mechanisms can contribute to its activation both at the transcriptional and post-translational levels [66]. The inhibition of cell surface PD-L1 upregulation during the UPR by either pharmacological or genetic means indicates that the IRE1α/XBP1 axis functions as a gatekeeper of PD-L1 expression in macrophages independently of IFNγ produced locally by T cells. By comparing gene expression in *Ern1*- and *Xbp1*-CKO macrophages it became apparent that *Ern1* but not *Xbp1* regulates a UPR-mediated PD-L1 gene expression.

Mouse studies showed that the sensitivity to PD-L1 blockade depends on PD-L1 expression in myeloid cells (macrophages and dendritic cells) and not on tumor cells [67, 68]. Remarkably, a recent report showed that ISRIB, a small molecule that reverses the effects of eIF2α phosphorylation downstream of PERK, reduces the abundance of the PD-L1 protein in murine *MYC^Tg^;KRAS^G12D^* liver cancer cells [51]. Whereas both reports agree on the role of the UPR in regulating PD-L1 expression, the discrepancy between the two studies creates an interesting conundrum as to why PD-L1 might be under the control of two different arms of the UPR in myeloid and tumor cells, respectively. Significantly, our analysis of tumor-infiltrating macrophages isolated from human endometrial and breast cancers indicates that the IRE1α gene signature is a better predictor of *CD274* (PD-L1) transcription than the PERK gene signature, confirming the conclusion reached in mouse macrophages, pointing to IRE1α as an important IFNγ-independent regulator of *CD274* in macrophages. Since PD-L1 serves as the ligand for PD-1^+^ T cells with exhausted [69] or regulatory phenotype [70], a plausible conclusion from the present study is that IRE1α inhibition in tumor-infiltrating myeloid cells could be used therapeutically to ameliorate the effects of immune dysregulation in the TME, including the downregulation of PD-L1, ultimately rescuing a failing immune surveillance and restoring immune competence locally.

A RIDD analysis in *Ern1* deficient macrophages showed a dramatic loss of the integrity and connectivity of RIDD genes compared to control (*Ern1 fl/fl)* macrophages. This provides initial mechanistic evidence that RIDD may be involved in shaping the immune landscape in the TME, including PD-L1 expression. A possibility is that upon IRE1α activation, RIDD degrades not only mRNAs but miRNAs as well, among which is miR-34a [71, 72], a miRNA also shown to target *CD274* (PD-L1) mRNA by directly binding to its *3*’- UTR [73, 74]. The loss of RIDD integrity shown here suggests that RIDD / miR-34a could represent the link between IRE1α and *CD247* gene expression. Future studies will need to address the role of RIDD in PD-L1-driven immune dysregulation in the TME.

In conclusion, we provide evidence in support of UPR-driven mechanisms as a source of immune dysregulation in the tumor microenvironment. We have identified the IRE1α/XBP1 axis as a critical signaling pathway in macrophage polarization to a mixed pro-inflammatory/immune suppressive phenotype, PD-L1 expression and tumor growth. Cell-nonautonomous IRE1α-dependent signaling has been proposed as a regulator of immune activation [75] and stress resistance and longevity in *C. elegans* [76], suggesting that the IRE1α/XBP1 axis may be central to intercellular communication during cellular stress. Here we further validate the view that UPR signals in the TME directly affect tumor-infiltrating macrophages promoting a complex immune dysregulation and defective tumor control *in vivo*. The fact that the IRE1α/XBP1 axis also regulate PD-L1 expression point to the UPR as a general mechanism for immune dysregulation at the tumor and immune cells interface with myeloid cells ultimately impairing the function of tumor specific T cells [28, 36] with loss of local immune surveillance.

## MATERIALS AND METHODS

### Cell lines and cell culture

Human cells lines colon carcinoma DLD1 and prostate PC3 and murine cell lines prostate TC1 and melanoma B16.F10 cancer cells were grown in RPMI or DMEM (Corning) supplemented with 10% FBS (HyClone) and 1% penicillin/streptomycin/L- glutamine, NEAA, sodium pyruvate, HEPES. All cells were maintained at 37°C incubation with 5% O_2_. All cell lines were mycoplasma free as determined PCR assay (Southern Biotech).

### Mice

APC mice were provided as a kind gift from Dr. Eyal Raz (UCSD). LysM. B6.129P2-Lyz2 ^tm1(cre)lfo^/J (LysM-Cre) mice were kindly provided by Dr. Richard Gallo (UCSD). ERN1^fl/fl^ and XBP1^fl/fl^ mice were kindly provided by Dr. Jonathan Lin (UCSD) who originally obtained them from Drs. Laurie Glimcher (Dana Farber, Harvard University) and Takao Iwawaki (RIKEN, Japan). All mice were housed in the UCSD vivarium according to approved protocols and animal welfare standards. Genotype of CKO mice were confirmed by PCR on tissue obtained by ear punch and digested according to a standard protocol.

### TERS Conditioned Medium (CM) Generation

DLD1 cells were induced to undergo ER stress through treatment of 300 nM thapsigargin (Tg) (Enzo Life Sciences) for 2 hours. Control cells were similarly treated with an equal volume of vehicle (0.02% ethanol). Cells were washed twice with Dulbecco’s PBS (Corning), and then incubated in fresh, standard growth medium for 16 hrs. Conditioned medium was then harvested, centrifuged for 10 min at 2,000 RPM, filtered through a 0.22-μm filter (Millipore), and treated to cells or stored at −80°C until use. For TERS priming, conditioned media was generated from homologous cell type unless otherwise specified. To measure IFNγ in TERS CM, QBeads (Intellicyt, Ann Arbor, MI) were used following manufacturer’s instructions. IFNγ was quantified on the iQue Screener PLUS (Intellicyt) using a standard cursive and manufacturer-provided template for analysis.

### BMDM and BMDC generation in culture

Bone marrow derived cells were procured by isolating the femur and tibia of specified host and flushing out the bone marrow using cold, unsupplemented RPMI growth media (Corning) using a 27 gauge needle and syringe. Hemolysis was performed using ACK Lysis buffer (Bio Whittaker). For macrophage differentiation, bone marrow cells were incubated one week in standard growth medium supplemented with 30% L929 conditioned medium (LCM) or m-CSF (origin) at concentration.

### ERAI activity assay

Cancer cell line reporter cells were transduced with the ERAI construct, originally described (234). Briefly, the pCAX-F-XBP1ΔDBD-venus (a kind gift from Dr. Iwawaki, Gunma University) underwent PCR using following primers: F: ctaccggactcagatctcgagccaccATGGACTACAAGGACGACG, R: gaattatctagagtcgcggccgcTTACTTGTACAGCTCGTCC. PCR fragments were cloned into pLVX-puro (Clontech) lentivirus vector with Gibson Assembly Mixture (NEB) according to manufacturer’s instruction. Stbl3 competent cells were transformed to produce the plasmid insert, whose presence was confirmed by sequencing. For production of lentivirus, 293FT (Invitrogen) cells were seeded in 10 cm dish and transfected with a plasmid mixture of ERAI plasmid and psPAX2 and pMD2G viral packaging plasmids. The supernatant of virus– producing transfected cells was collected every 24 hrs for three days post transfection. Viral supernatant was concentrated by 10% PEG-8000 and pelleted with 2000 x g for 40 min at 4C and re-suspended PBS. Target cancer cells were transduced with lentivirus by adding supplementing with polybrene (8 μg/mL) to virus containing solution and loaded onto B16.F10 cancer cell line. Lines were transduced for 48 hours. Following, cells were washed twice with PBS and positively selected for using puromycin (2 μg/mL) for two weeks. In some instances, positively transduced cells were then stimulated for Venus expression and were sorted by FACS (BD) to isolate high expressing clones. Lines were maintained under puromycin.

### Flow cytometry

Single cell suspensions of myeloid cells were separated and stained for CD80 (B7-1) (BD Biosciences), PD-L1 (CD274) (BD Biosciences), and CD86 (BD Biosciences). Viable cells were determined by 7AAD exclusion and data were acquired using a FACScalibur flow cytometer (BD). Flow results were analyzed using CellQuest Pro (BD) and Flow JO (Tree Star) software.

### RT-qPCR

mRNA was harvested from cells using Nucleopsin II Kit (Machery-Nagel) or enzymatically using the Zygem RNAgem Tissue PLUS kit (Microgembio, New Zealand). Concentration and purity of RNA was quantified the NanoDrop (ND-1000) spectrophotometer (Thermo Scientific) and analyzed with NanoDrop Software v3.8.0. RNA was normalized between conditions and cDNA generated using the High Capacity cDNA Synthesis kit (Life Technologies). RT-qPCR was performed on ABI 7300 Real-Time PCR system using TaqMan reagents for 50 cycles using universal cycling conditions. Cycling conditions followed manufacturer’s specifications (KAPA Biosystems). Target gene expression was normalized to *β-actin* and relative expression determined by using the –ΔΔCt relative quantification method. Primers for qRT-PCR were purchased from Life Technologies: Arg1, (Mm00475988_m1), Cd274 (Mm03048248_m1), Chop (Mm00492097_m1), Grp78 (Mm00517691_m1), Il6 (Mm99999064_m1), Il23-p19 (Mm00518984_m1), and Tapbp (Mm00493417_m1).

### Western Blot Analysis

After treatment, cells were washed with ice cold PBS and suspended in the RIPA Lysis Buffer system: 1X RIPA buffer and cocktail of protease inhibitors (Santa Cruz Biotechnology). Cell lysates were centrifuged at 16,000g for 15 min and the supernatants were extracted. Protein concentration was determined using Pierce BCA Protein Assay Kit (Thermo Scientific). Samples were heat denatured and equal concentrations of protein were electrophoresed on 4-20% Mini-PROTEAN TGX Precast Gels (Bio-Rad) and transferred onto 0.2 µm PVDF membrane in Tris-Glycine transfer buffer containing 20 % methanol. The membranes were blocked with 5% non-fat milk in TBS containing 0.1 % Tween-20 (TBS-T) for 1 h at room temperature, and subsequently incubated with diluted primary antibodies overnight at 4 °C. Membranes were washed for 5 min at room temperature 3 times by TBS-T, incubated with secondary antibody conjugated with horse radish peroxidase (HRP) in 5 % non-fat milk for 1 h at room temperature, and washed for 5 min at room temperature 3 times by TBS-T. Immuno-reactivity was detected by chemi-luminescence reaction using Pierce ECL Blotting Substrate (Thermo Scientific). Primary antibodies used were: rabbit monoclonal antibody to IRE1α (clone 14C10) (Cell Signaling Technology), rabbit polyclonal antibody to XBP-1s (#83418) (Cell Signaling Technology), goat polyclonal antibody to GAPDH (A-14) (Santa Cruz Biotechnology). Bound primary antibodies were revealed by the following secondary antibodies: HRP-conjugated goat antibody to rabbit IgG (Cell Signaling Technology), and HRP-conjugated donkey antibody to goat IgG (sc2020) (Santa Cruz Biotechnology).

### Tumor studies

For orthotropic tumor implantation model, B16.F10 cancer cells (n=4) were detached from plastic, washed twice with cold PBS, and resuspended at a concentration of 3e5cells/ml in PSB. Host C57BL/6 or transgenic ERAI mice (a kind gift from Dr. T. Iwawaki (Gunma University)) were subcutaneously injected with 100 μl (3e4 cells) of cell suspension into the right hind flank. After approximately 22 days, mice bearing tumors greater than 1 cm were sacrificed. For tumor growth studies, B16.F10 were subcutaneously injected in C57BL/6 (WT) or TLR4 KO mice (a kind gift from Dr. M. Corr (UCSD)). Tumor establishment was first determined by palpation and size was then measured in two dimensions using calipers. When tumors reached > 20 mm in any one dimension or after 30 days post implantation, whichever came first, mice were sacrificed. Tumor volume was calculated using the ellipsoid volume formula, V = 1⁄2 (H x W^2^). All mice were sacrificed when any tumor reached 20 mm in any one dimension, per UCSD animal welfare standards, or after 30 days post implantation. Tumor volume was calculated using the ellipsoid formula: V = ½ (H x W^2^).

### Isolation of CD11b^+^ and F4/80 cells

For B16.F10 model: B16.F10 cancer cells (n=5) were subcutaneously injected (3e4) into the right hind flank of C57BL/6 mice. After approximately 22 days, mice bearing tumors greater than 1 cm were sacrificed. For APC model: APC mice were genotyped for *APC* mutation to confirmed homozygosity of transgene. At approximately 12-15 weeks of age, APC mice were sacrificed by cervical dislocation. The small intestine was removed from host and cut longitudinally, running parallel to the intestinal lining. Adenomas lining the intestine were excised using an open blade and pooled, respective to the host, in ice cold PBS supplemented with 0.5% (w/v) bovine serum albumin (BSA). For both model systems: once the tumor, spleen, and bone marrow were isolated from tumor bearing hosts, tissues were dissociated through enzymatic digestion (TrypLE) at 37°C for 30 min on a rocker 85 plate, followed by cell straining through a 22 μm filter in ice cold PBS + 0.5% (w/v) BSA. Cell suspensions were then stained for CD11b^+^ positivity by first using a CD11b-biotin conjugated antibody (BD Biosciences) and incubated for 15 min at 4°C. Cells were then washed twice with PBS + 0.5% BSA and positively selected by magnetic separation using a biotin isolation kit (Stem Cell) according to manufacturer’s specifications. F4/80^+^ macrophages were isolated from subcutaneous B16.ERAI tumors from the right hind flank *Ern1 x LysMCre* or *fl/fl* mice. After approximately 22 days, mice bearing tumors > 1 cm in length were sacrificed. Tumors and spleens were isolated, tissues were dissociated through enzymatic digestion (TrypLE) at 37°C for 30 min on a rocker 85 plate, followed by cell straining through a 22 μm filter in ice cold PBS + 0.5% (w/v) BSA. Cell suspensions were then stained for F4/80^+^ positivity by first using a F4/80-PE conjugated antibody (StemCell Technologies Cat# 60027PE.1) and incubated for 15 min at 4°C. Cells were then washed twice with PBS 0.5% BSA and positively-selected by magnetic separation using PE Positive Selection Kit II (StemCell Technologies) according to manufacturer’s specifications.

### RNASeq analysis

RNA was extracted from wild type or Ern1 CKO BMDM that were untreated or treated with TERS CM for 18 hours using the Nucelospin RNA kit (Macherey Nagel). Each group consisted of 3 independently-derived BMDM. RNA sample purity was ascertained by the Nanodrop quantification method. Single end stranded RNA libraries for were sequenced on an Illumina HiSeq 4000. All samples and replicates were sequenced together on the same run. All 12 mouse RNA-seq transcript quantification was performed with sailfish version 0.9.2 [77], using the GRCm38 mouse transcriptome downloaded from Ensembl (URL: ftp://ftp.ensembl.org/pub/release-97/fasta/mus_musculus/cdna/Mus_musculus.GRCm38.cdna.all.fa.gz) with default parameters. The 33 RIDD target genes were collected from [49]. We z-scored these RIDD target genes within each group separately (*Ern1* fl/fl and *Ern1* CKO) and then mean value was calculated and compared between different phenotype (untreated vs TERS CM treated) within each group.

### Ordinary Least Squares (OLS) linear model predicting PD-L1 using IRE1α pathway and PERK pathway downstream genes

OLS models were fitted and compared using the python (version 2.7.15) statsmodels package (version 0.9.0). We collected IRE1α pathway (R-HSA-381070.1) and PERK pathway (R-HSA-381042.1) downstream genes from REACTOME [54]. Each gene was z-scored to ensure a mean of 0 and standard deviation of 1. Because quantification of transcript levels is noisier when genes are expressed at low levels, we implemented a filter to remove genes expressed under a certain threshold and evaluated pathway scores at thresholds ranging from 0 to 1000 reads. We then fitted models at different thresholds to evaluate robustness of the model to choice of threshold. Models were fitted using the formula:

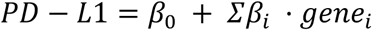

Nested OLS models with ERN1 only and ERN1 + PERK were compared using the Aikake information criterion (AIC). For each model, the AIC was calculated as AIC = 2*k* - 2ln(*L*), where *k* represents the number of estimated parameters, and *L* represents the likelihood function for the model. Models were compared using the formula exp((AIC_min_ − AIC_i_)/2), which represents the relative likelihood of model i with respect to the best available model.

### Statistical analysis

To determine if differences between groups were statistically significant for PCR experiments, groups were compared using unpaired student’s *t*-tests with Welch’s correction. Statistically significant differences are indicated as follows: *p<0.05, **p<0.01, ***p<0.001, ****p<0.0001. Statistical significance in tumor growth experiments was determined using the Mann-Whitney *t* test and survival curves were generated by the Kaplan-Meier method.

## Acknowledgements

This work was supported in part by grant RO1 CA220009 to M.Z. and H.C. J.J.R. acknowledges the support of the Frank H. and Eva B. Buck Foundation. The authors thank Valentina Ferrari for performing the QBeads assay.

## Author Contributions

Conceptualization, M.Z., J.J.R., H.C., Reagents and Specimens, T.I., J.L., Data Collection, A.B., J.J.R., S.X., S.S., A.L., G.A., K.J., Manuscript writing, M.Z., S.X, Manuscript revisions, J.J.R., S.S., J.L., H.C., Supervision, J.J.R., H.C., M.Z. Funding acquisition, M.Z., H.C., J.J.R.

